# A bHLH transcription factor (VfTT8) underlies *zt2, the locus determining zero tannin content in faba bean (Vicia faba L.)*

**DOI:** 10.1101/2020.03.06.980425

**Authors:** Natalia Gutierrez, Carmen M. Avila, Ana M. Torres

## Abstract

Faba bean (*Vicia faba* L.) is an important protein-rich fodder crop widely cultivated in temperate areas. However, antinutritional compounds such as condensed tannins, limit the use of this protein source in monogastric feed formulations. Previous studies demonstrated that two recessive and complementary genes, *zt1* and *zt2*, control the absence of tannin and the white flower colour in faba bean. An ortholog of the *Medicago* WD40 transcription factor, (*TTG1*) was reported to encode the zt1 phenotypes but the responsible gene for zt2 is still unknown. A candidate gene approach combined with linkage mapping, comparative genomics and gene expression has been used in this study to fine map the *zt2* genomic region and to identify the regulatory gene controlling both traits. Seventy-two genes, including 23 regulatory genes (MYB and bHLH) predicted to be associated with anthocyanin expression together with WRKY proteins were screened and genotyped in three mapping populations. The linkage groups constructed identified the regulatory gene, *TRANSPARENT TESTA8* (*TT8*), encoding a basic helix-loop-helix transcription factor (bHLH), as the best candidate for *zt2*. This finding was supported by qPCR analyses and further validated in different genetic backgrounds. Accordingly, VfTT8 was down-regulated in white flowered types, in contrast to the levels of expression in wild genotypes. Our results provide new insights on the regulatory mechanisms for tannin biosynthesis in faba bean and will favour the development of an ultimate zt2 diagnostic marker for the fast generation of new value-added cultivars free of tannins and improved nutritional value.

## Introduction

Since the beginning of the agriculture legume species with high levels of protein in seeds and foliage have been essential sources of dietary protein for humans and animals. Faba bean (*Vicia faba* L.) is a worldwide cultivated grain legume, known for its great potential for yield and high protein contents varying from 25-40%^1^. Nevertheless, the presence of tannins, primarily located in the seed coat^2,3^, limit successful deployment of this protein rich fodder crop in feedstuffs for monogastric animals. These polyphenolic compounds are present in the hull portion of the seeds at different concentrations depending on variety, maturity, location and growth conditions^4^. Tannins play an important role in plant growth and reproduction, providing protection against biotic and abiotic stresses^5^. Despite these defensive effects, they have a negative impact on nutritive value and digestibility being responsible for decreases in feed intake, growth rate, feed efficiency, net metabolizable energy and protein digestibility in experimental animals^6^. Low-tannin content generally results in higher protein and energy digestibility for monogastric animals^7^, consequently, the development of faba bean tannin-free cultivars is a key breeding objective to enhance the nutritional quality and broaden its use in livestock feed industry.

Flavonoids are the most common group of polyphenolic compounds in the plant kingdom^8^. Thanks to the easily detectable mutant phenotypes in flower and seed pigmentation, the flavonoid biosynthesis is one of the best-studied secondary metabolic pathways. These compounds play important roles in the interactions of plants with their environments serving as pigments, signalling molecules, protectants against biotic and abiotic stresses and UV protection^9,10^.

Proanthocyanidins (condensed tannins) and anthocyanins are major flavonoid end-products of a well conserved family of aromatic molecules and both compounds are produced by related branches of the flavonoid pathway and utilize the same metabolic intermediates. Two classes of genes are required for anthocyanin and tannins biosynthesis, the structural genes encoding the enzymes that directly participate in the formation of different flavonoids, and the regulatory genes that control the transcription of structural genes^11^. Mutations in either structural or regulatory genes have been shown to lead to a loss of pigmentation in several plant species^12–15^.

Many of the structural and regulatory genes have been identified and cloned in model plants including Arabidopsis, maize (*Zea mays*), snapdragon (*Antirrhinum majus*) and Petunia^9,16–18^. The common structural genes in this pathway are chalcone synthase (CHS) chalcone isomerase (CHI), flavanone 3-hydroxylase (F3H) and dihydroflavonol 4-reductase (DFR) which are highly conserved among plant species. Likewise DNA-binding R2R3-MYB transcription factors, basic-Helix-Loop-Helix (bHLH) transcription factors, and WD40 repeat proteins are known to form MYB-bHLH-WD repeat (MBW) complexes, which activates the transcription of structural genes in the anthocyanin pathway^19^.

In previous studies, we checked whether the faba bean genes producing white flower and low tannin phenotypes were in the anthocyanin biosynthetic pathway. Thus, we amplified the key common structural genes (CHS, CHI, F3H and DFR), in order to identify polymorphisms that might account for functional gene changes. Different bioinformatics tools, including BLAST and multiple sequence alignment were used to design primers suitable for their amplification in contrasted faba bean populations. However neither the subsequent linkage analysis nor the expression studies by RT-PCR pointed to any of these genes as possible regulators of the zt-1 and zt-2 phenotypes^20^.

In pea, the absence of pigmentation in the flower is the result of mutations in loci *A* and *A2* encoding a bHLH transcription factor and a WD40 protein, respectively^21^. In lentils the zero-tannin trait is controlled by a single recessive gene (*tan*) that encodes a bHLH transcription factor, homologous to the *A* gene in pea^22^. In faba bean, the zero-tannin (*zt*) trait is a simple character, governed independently by two complementary recessive genes *zt1* and *zt2*^2,23^. These genes interrupt the synthesis of anthocyanins or of their precursors at different steps in this pathway, giving plants with white flowers and no tannin content^3,24^. Considering the partial outcrossing behaviour of faba bean, crosses between zt1 and zt2 individuals will produce coloured plants with tannins that might contaminate the crop. For this reason, it is also important to know which gene is present in each tannin-free accession in order to select the appropriate genitors in breeding programs.

Recently, an ortholog of the *Medicago* WD40 transcription factors, *Transparent Testa Glabra 1* (*TTG1*) in the faba bean chromosome (chr.) II, has been reported as the gene encoding the zt1 phenotypes^25–28^. Likewise, the locus *zt2* was found to be in the distal part of the chr. III^26,28,29^ but the responsible gene is still unknown. After this initial finding, the later authors^29^ reported a KASP (Kompetitive Allele-specific PCR) marker (SNP marker Vf_Mt7g100500_001), located 10.5 cM apart from the flower colour (*zt2* locus), as a reliable marker to discriminate low-tannin faba bean plants carrying *zt2*. Considering the relatively high genetic distance reported, uncertainty and false selection may occur due to recombination events. To exclude such recombination, marker enrichment of the genomic region is needed to identify the underlying gene and further facilitate a reliable marker-assisted selection in breeding programs.

The main goals of the present study were, therefore, to saturate the *zt2* genomic region and to identify putative candidates for the white flower trait. To do so, we have used a combination of genetic linkage, association studies and comparative genomic approaches with the legume model *Medicago truncatula* (http://www.medicagohapmap.org/fgb2/gbrowse/mt35/). As mentioned above, the first step was the assignment of the *zt2* gene to a specific faba bean chromosome using a linkage approach. Markers flanking *zt2*^30^ *together with other Medicago* genes were genotyped in three RIL populations used to develop a composite map of this species^31^. The approach allows us to assign the target loci to the distal part of the chr. III^26,28^. Upon confirmation of the chromosomal region containing the flower colour locus, the next step was to exploit the extensive collinearity with *Medicago* to saturate the target region and mine candidate genes associated with the trait. The approach pinpointed a number of regulatory genes (MYB and bHLH) predicted to be associated with the anthocyanin expression. Once the gene sequences were available our objective was to associate the mutations with the target phenotypes using different mapping populations and genetic backgrounds.

In summary, in the present study we describe a detailed approach for fine mapping, identification and validation of the gene underlying *zt2* in faba bean. This outcome will facilitate the development of a diagnostic marker, fully efficient for the selection of zt2 cultivars free of anti-nutritional compounds.

## Material and Methods

### Plant material and sample collection

Three mapping populations (two F_2_ and one F_7_) segregating for tannin content (TC) and flower colour (FC), wild spotted type vs. white flower, were used in this study: 1) M x D: 50 F_2_ individuals originated from the cross of MAYA (M), with spotted flowers and high TC and DISCO (D), a white flower/zero tannin inbred line, carrying the *zt2* gene, 2) W x D: 56 F_2_ individuals derived from the cross of WIZARD (W), a wild-type flower line with high TC and DISCO (D) and, 3) Vf6 x zt2: 62 F_7_ individuals, where the maternal parent (Vf6) has spotted flowers and high TC while zt2 is a tannin-free white flowered line. The populations were selfed at the IFAPA of Córdoba, Spain. Plants were grown in insect proof cages to avoid outcrossing. Young leaves were collected from all samples and DNA was isolated according ^32^. The concentration and purity of DNA was measured by a NanoDrop ND-1000 spectrophotometer. The final concentrations of all DNA samples were adjusted to 10 ng/μl for high-throughput genotyping.

### Phenotypic evaluation

F_2_ plants from crosses M x D and W x D were selfed to the F_3_ generation and the FC was recorded at each F_3_ family (15 plants per line) to infer their corresponding genotype (homozygous or heterozygous). In the RIL population (Vf6 x zt2), FC was scored in all the lines self-pollinated from F_2_ to F_7_. Segregation ratios were tested against the expected ratio using a chi-square analysis for goodness of fit. Phenotypic data on FC in the three populations are available in the Table 1.

**Table 1.** Segregation of flower colour (FC) in the three populations used in this study. The observed and expected ratios correspond to the homozygous wild type: heterozygous: homozygous white flower colour in F_2:3_ generations and homozygous wild type: homozygous white flower colour in F_7_ RIL population. **\* α** = 0.5

| Population | Generation | Individuals evaluated | Observed Ratio | Expected Ratio | $\chi^2$ | p-Value * |
| --- | --- | --- | --- | --- | --- | --- |
| M x D | F<sub>2:3</sub> | 50 | 12:30:8 | 1:2:1 | 2.64 | 0.27 |
| W x D | F<sub>2:3</sub> | 56 | 18:24:14 | 1:2:1 | 1.71 | 0.42 |
| Vf6 x zt2 | F<sub>7</sub> | 62 | 36:26 | 1:1 | 1.61 | 0.20 |

### Genomic location and saturation methodology

RAPD markers flanking *zt2*^30^, *together with a number of ESTs from Medicago* and *Pisum* developed within the Grain Legumes Integrated Project (GLIP-Food-CT-2004-506223), were genotyped in different faba bean RIL (Recombinant Inbred Lines) populations and further used to develop a consensus map in this species^31^. The approach ascribed the target *zt2* region in the distal part of faba bean chr. III between the ESTs markers Pis-Gen-8-1-1 and LG38^26,28^. Using the sequence information deposited in the *M. truncatula* Genome Database (MTGD) (http://www.medicagogenome.org/), these genes corresponded to Medtr1g040675 and Medtr1g087900, respectively, spanning 24.3 Mb.

For marker saturation and candidate gene identification we performed a detailed genomic analysis of this syntenic region. Seventy-two genes, including regulatory genes (MYB and bHLH) predicted to be associated with anthocyanin expression together with WRKY proteins were selected to be screened for polymorphism in the three mapping populations (Supplementary Table S1).

### Sequencing and genotyping technique

The 72 genes selected include six ESTs markers previously assigned to the target interval^26,31^ and 23 newly designed *M. truncatula* markers associated with the anthocyanin expression. The remaining 42 genes were genotyped using the Kompetitive Allele-Specific PCR (KASPar) assays developed by Webb^27^ and provided by LGC genomics (https://www.biosearchtech.com/products/pcr-kits-and-reagents/genotyping-assays/kasp-genotyping-chemistry) (Supplementary Table S1).

Different tools, including BLASTn and multiple sequence alignment analysis were used to design primers suitable for DNA amplification. The sequences were BLASTed against an in-house faba bean transcriptome^33^ to obtain the orthologous sequence in faba bean. For intron-exon boundary prediction and primer design, alignment between sequences were implemented in Geneious v.7.1.5 (https://www.geneious.com) The information about primer sequence, annealing temperature, marker type, orthologous sequence in faba bean, locus name and annotation in *M. truncatula* of the 23 new markers are available in Supplementary Table S2.

To obtain the sequences from DNA samples (parental lines and two contrasting individuals from each population), PCR reactions were conducted in a total volume of 25 µl, using 5 µl DNA of each sample, 200 nM of each primer, 2 mM MgCl2, 200 µM dNTP and 0,6 U Taq polymerase Biotools (B&M Labs, S.A., Madrid, Spain). Amplification products were purified using a standard protocol for DNA precipitation with sodium acetate and ethanol (1/10 3M sodium acetate, 2 v/v ethanol) (https://www.thermofisher.com/es/es/home/references/protocols/nucleic-acid-purification-and-analysis/dna-protocol/sodium-acetate-precipitation-of-small-nucleic-acids.html#comergent_product_list_15878). PCR products were sequenced by Sanger at STABVIDA (Caparica, Portugal). For each sample, four PCRs were done, and the repeated products were then mixed for sense and antisense strand sequencing.

Sequence analysis and identification of polymorphisms were conducted using Geneious v.7.1.5. For each gene, the sense and antisense sequences were aligned and the consensus sequence further analyzed with BLASTn against the *M. truncatula* genome (Mt4.0) to assign the percent-consensus-identity. Sequence polymorphisms between the samples were transformed to CAPS (Cleaved Amplified Polymorphism) markers by restriction enzyme digestion of the PCR products. The digestions were carried out in a 15 µl volume of PCR using 5 µl of the amplified products and 3U of the respective restriction endonuclease (ThermoFisher Scientific). The digestions were incubated overnight at their respective temperature. The restricted fragments were separated in 2% agarose gel. Detailed information of the specific gene-based markers designed in this study together with the KASPar markers assayed, is shown in the Supplementary Table S1.

### Linkage maps calculation

The segregation of each marker was tested for goodness of fit to the expected segregation ratio (1:2:1 in the two F_2_s families and 1:1 ratio in the RIL population), using the chi-square test. Linkage maps were constructed using JoinMap 4.1^34^. Markers were grouped using regression mapping algorithm option at a minimum LOD score threshold of 4 and maximum recombination frequency of 0.4 as general linkage criteria to establish linkage groups (LG). Recombination fractions were converted to centimorgans (cM) using the mapping function of Kosambi^35^.

### qPCR samples, RNA isolation, cDNA synthesis and primer design

The most important guidelines of the MIQE checklist^36^ were considered according to the practical approach for quantitative real-time PCR (qPCR) experiments proposed by Taylor^37^. The expression profile of candidate genes was analysed in pigmented and white flowers from the Vf6 x zt2 and M x D populations at two different developmental stages. Stage 1 (S1) included immature flowers buds of approximately 1,5 cm length while the stage 2 (S2) included young flowers of 2 cm length and black colour apparent at the top of the petal in the wild types. CYP2 and ELF1A, previously reported as the most stable genes for normalization of the gene expression in the tannin content experiment were used as reference^38^.

Only petal tissue was used for total RNA extraction. In order to minimise variation in gene expression among individual plants and stages, petal tissue from three individuals were collected and pooled together for RNA isolation. Finally, three pools of biological replicates were used per individual genotypes. Each sample was frozen in liquid nitrogen and stored at −80°C until RNA extraction. In short, the experimental design consisted in a total of 24 samples (2 genotype × 2 stages of development × 2 technical repetitions × 3 biological repetitions).

The Direct-zol RNA Kit (Zymo Research) was used to isolate a high-quality RNA directly from samples in TRIzol Reagent®. RNA concentration was determined by measuring the optical density using a NanoDrop spectrophotometer. Only the RNA samples with A_260_/A_280_ ratio between 1.9 and 2.1 and A_260_/A_230_ greater than 2.0 were used in the analysis. cDNA of each sample was obtained by iScriptTM cDNA Synthesis kit (Bio-Rad, Hercules, California) and diluted to a concentration of 20 ng/µL. A pooled sample comprising all samples considered in the experiment was included for each gene as inter-run calibrator to detect and correct inter-run variation. No-template controls were also included.

Specific primer pairs were designed using the following criteria: Tm of 60 ± 1°C and PCR amplicon lengths of 80–120 bp, yielding primer sequences with lengths of 19–26 nucleotides and GC contents of 40–80%. Designed primers were synthesized by Integrated DNA Technologies (Leuven, Belgium). Detailed information on the primers designed for RT-qPCR is shown in Table 2.

**Table 2.**
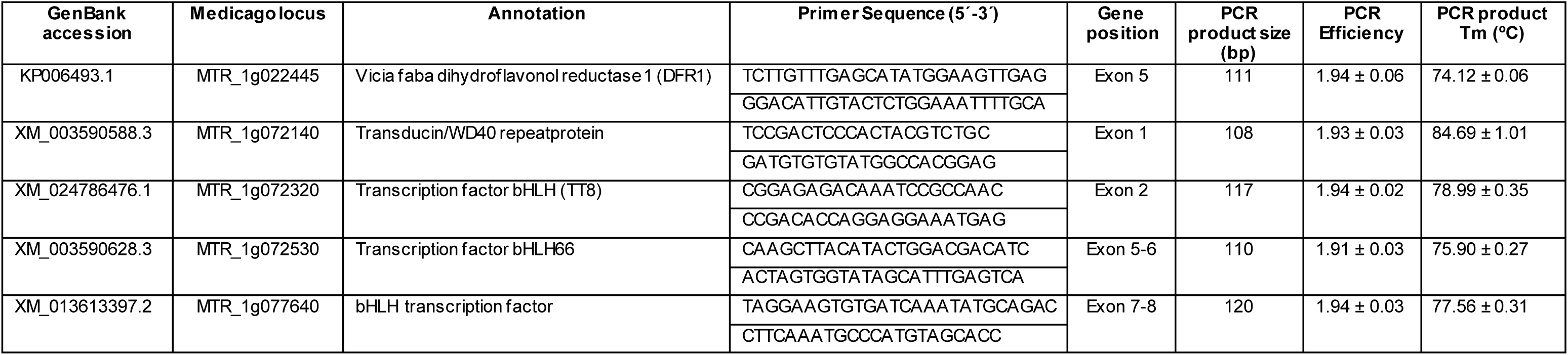
Information on the primers for RT-qPCR designed in the study. GenBank accession, medicago locus, annotation NCBI, primer sequences, gene position and amplicon size. Primer PCR efficiency and PCR product Tm data represent mean values ± sd.

### Real time qPCR assays

The qPCR was carried out using the iTaqTM Universal SYBR® Green Supermix on an ABI PRISM 7500 Real Time PCR System (Applied Biosystems, Foster City, CA, USA). Experiments were performed in 96-well optical reaction plates (P/N 4306737) with MicroAmp optical adhesive film (P/N 4311971) from Applied Biosystems. A master mix with total volume of 11 µL for each PCR run was prepared, containing 5 µL of diluted cDNA (10 ng/µL), 5 µL of iTaqTM Universal SYBR® Green Supermix (Bio-Rad, Hercules, California) and a primer pair with a concentration of 0.22 µM each. The PCR conditions were 95°C for 10 min followed by 40 cycles at 95°C for 15 s and 60°C for 1 min. The specificity of the of the PCR products was confirmed by single and sharp peaks at the melting curves analysis after 40 amplification cycles.

Fluorescence was analyzed using 7500 Software v2.0.1. All amplification plots were analyzed using a threshold values of 0.2 to obtain Cq (quantification cycle) values for each gene-cDNA combination. PCR efficiency of each primer pair was determined for all samples by LinRegPCR program v.11 ^39^, using raw normalized (Rn) fluorescence as input data.

The relative gene expression (RGE) was calculated using the advanced quantification model with efficiency correction, multiple reference genes normalization and use of error propagation rules described by Hellemans^40^ (equation 1), where RQ = E^ΔCt^, being E = PCR efficiency for each primer and Ct = the number of cycles needed to reach 0.2 arbitrary units of fluorescence. The two reference genes (RG) used for data normalization were CYP2 and ELF1A ^38^.

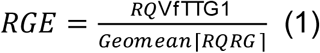

The RGE values were log-transformed and the significance values to determine differences of the expression of genes were obtained by the ANOVA test using the R programming language.

## Results

### Segregation analysis of flower colour (FC)

Of the 50 F_2:3_ plants screened from the M x D population, 12 had spotted flower, 30 were heterozygous/spotted and 8 showed white flower, fitting a 1:2:1 ratio (χ^2^ = 2.64; p = 0.27). Further, the 56 F_2:3_ families from cross W x D, also exhibited a good fit for the expected 1:2:1 (spotted: segregating: white) ratio (χ^2^ = 1.71; p = 0.42). Similarly, the 62 RILs evaluated fit the 1:1 (spotted:white) ratio (χ2 = 1.61; p = 0.20) (Table 1). As expected, results from FC segregation in these populations confirmed the monogenic control of the gene with a dominant *ZT2* (spotted flower) and a recessive *zt2* (white flower) allele.

### Genotyping and linkage analysis

The collinearity between faba bean and the model *Medicago* was exploited to develop new markers and fine map *zt2,* in the distal part of faba bean chr. III. Seventy-two genes markers, including genes encoding MYB, bHLH and WRKY proteins, were surveyed in the interval flanked by Medtr1g040675 and Medtr1g087900 (Supplementary Table S1). Six of the markers were EST previously used by Satovic^31^. Moreover, we designed 23 new primer pairs in genes putatively associated with the anthocyanin expression while the remaining 43 genes, were genotyped using the KASPar technique.

Thirty out of the 72 genes tested revealed polymorphism in at least in one of the three faba bean populations (23 KASP-SNPs, 6 CAPs, 1 Amplified Length Polymorphism/ALP), 12 of them being anchor markers, common among populations (Supplementary Table S1). Only five of the 23 candidate genes predicted to be associated with anthocyanin expression resulted polymorphic and were further genotyped and mapped. The most polymorphic population was M x D with 20 markers, followed by Vf6 x zt2 (15 markers) and W x D (13 markers).

Three linkage groups were constructed (Fig. 1). Fine mapping analysis showed that in cross M x D, *zt2* is localized in a window of 6.1 cM between markers Vf_Mt1g072140 and Vf_Mt1g072320 whereas in W x D, *zt2* is in a window of 8 cM between Vf_Mt1g072320 and GLPSNP. Finally, in the RIL population (Vf6 x zt2) the target gene was flanked by Vf_Mt1g072320 and Vf_Mt1g077640 (Figure 1). Collinearity between faba bean chr. III and *M. truncatula* chr. I was high except for two markers Vf_Mt7g035110 and Vf_Mt7g100500 that revealed some chromosomal rearrangements, due to insertions from Mt Chr. 7. All markers maintained a similar mapping order in the three populations being Vf_Mt1g072320, a TT8 transcription factor, which we will henceforth refer to as VfTT8, the closest to FC in the three populations (2,8 cM; 1,1 cM and 0.8 cM, respectively). These outcomes pointed to *TRANSPARENT TESTA8*, encoding a bHLH transcription factor (Nesi et al., 2000), as the best candidate for zt2 in *Vicia faba*.

**Fig. 1.**
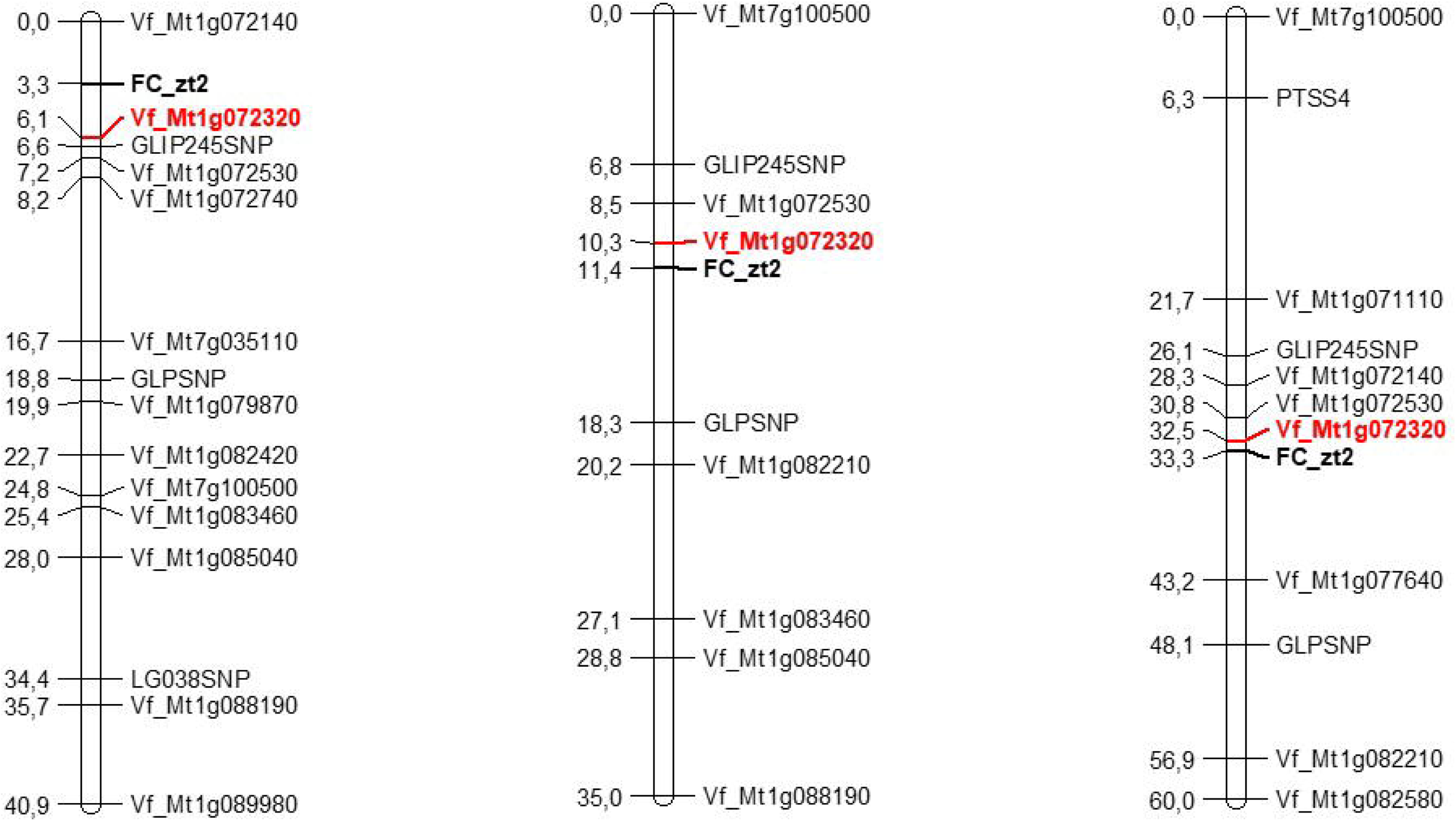
Linkage maps from three faba bean populations segregating for flower colour and carrying the gene zt2. M x D F_2_ population (left), W x D F_2_ population (middle) and Vf6 x zt2 RIL population (right). Markers names are on the right and the estimated map distances are shown on the left. Recombinant fractions were converted to centiMorgans using the mapping function of Kosambi.

### Haplotype analysis

In the stage of fine mapping, the 168 individuals (50 F_2:3_ from MxD, 56 F_2:3_ from W x D and 62 RILs from cross Vf6 x zt2), were screened to identify recombinants among flanking markers (Supplementary Table S3). Two individuals (in W x D and Vf6 x zt2) or three (in M x D), were found to have at least one recombination event between *zt2* and the closest marker Vf_Mt1g072320. A number of crossovers, ranging from 5 to 9, were also observed in the rest of flanking markers and crosses except for Vf_Mt7g100500, reported by Zanotto^29^ as a reliable marker for the selection of *zt2.* Vf_Mt7g100500 revealed, however, the highest number of misclassified individuals in the three populations (17, 10 and 20, respectively), contradicting the previous report and precluding the reliable use of this marker in faba bean breeding programs. According to these recombination events, Vf_Mt1g072320 appears to be the best candidate in discriminating low tannin individuals carrying the *zt2* gene.

Only in case of Vf_Mt1g072320, were designed two primers pairs in different genomic regions with the aim of ensuring the amplification using different genetic backgrounds and achieve the best resolution in the corresponding banding patterns (Supplementary Table S2). Thus, using primer pair 1, the recessive allele linked with the *zt2* locus in crosses W x D and Vf6 x zt2 is *G.* Therefore, considering identity by descent in the germplasm analyzed and without recombination between the marker and the *zt2* locus, white flower genotypes are expected to be *GG* whereas the wild flower genotypes are expected to be *AG* or *AA.* The primer pair 2 was only used in cross M x D and the corresponding genotypes were CC in white flower zt2 genotypes and *TC* or *TT* in wild individuals (Supplementary Table S3).

### Expression Analysis of candidate genes

With the objective to corroborate VfTT8 as the *zt2* gene, qPCR analysis was performed on flower tissues of two individuals from Vf6 x zt2 RILs, contrasted in FC. The candidate genes assayed were the most closely linked to FC_*zt2* in the three linkage maps (Fig 1): MTR_1g022445, a DFR1 gene (reported by Ray^41^), MTR_1g072140 a WD40 protein and MTR_1g07230, MTR_1g072530 and MTR_1g077640, all being bHLH transcription factors. cDNA from RIL6 (with pigmented flower) and RIL2 (white flower) at two different developmental stages (S1 and S2) were analysed.

To verify the sensitivity and specificity of the qPCR, we tested the presence of genomic DNA (gDNA) contamination in the cDNA samples, using the primer pairs for MTR_1g072530 and MTR_1g077640, which span an exon-exon junction (Table 2). The first primer pair amplified a product with 195 bp using gDNA as template whereas the band was of 110 bp using cDNA as template. In case of the primer pair MTR_1g077640, the amplified band was of 224 bp or 120 bp using gDNA or cDNA as template, respectively. None of the cDNAs tested revealed bands corresponding to residual genomic DNA, thus indicating the purity of samples. The dissociation curve analysis confirmed that all the primer pairs used, produced a single and specific PCR product (Table 2). The mean PCR efficiency for all genes was up to 95.5%, while for the reference genes (CYP2 and ELF1A) the mean PCR efficiency was 1.88 ± 0.009 and 1.88 ± 0.01 (mean ± sd), respectively.

The overall expression pattern of the MTR_1g072320 in both flower stages was found significantly higher (p < 0.001) in the V6 x zt2 wild-type line (RIL6) when compared to the RIL2, being strongly down-regulated in this white flowered genotype (Table 3 and Fig. 2). To validate these results, the expression of MTR_1g072320 was assayed in a second faba bean population (M x D). We used cDNA from two F2 individuals with pigmented flower (G1 and G11) and white flower (G5 and G29). The expression pattern of MTR_1g072320 matched to that obtained in the Vf6 x zt2 RILs, showing highly significant differences in the expression level (p < 0.001) of contrasted FC genotypes. Our results confirmed a strong down-regulation of the MTR_1g072320 expression in white flowered genotypes. These outcomes further support that VfTT8 is the gene responsible for the loss of flower pigmentation and the absence of tannins in the *zt2* genotypes.

**Table 3.** Gene expression ratios. Values indicate log average expression ratios of three biological replicates from each of the genotypes used in the study. RIL6 (wild type) and RIL2 (mutant type) are F_7_ individuals from the Vf6 x zt2 population. Statistically significant regulation (p < 0.001), is indicated in bold.

| Medicago locus | RIL population Vf6 x zt2 |  |  |  |
| --- | --- | --- | --- | --- |
|  | RIL6 (wild type) |  | RIL2 (zt2) |  |
|  | Stage 1 | Stage 2 | Stage 1 | Stage 2 |
| MTR_1g022445 | 0,08 | 0,28 | -0,18 | -0,19 |
| MTR_1g072140 | -0,49 | -0,09 | -0,42 | -0,16 |
| MTR_1g072320 | **0,47** | **0,81** | **-0,64** | **-0,64** |
| MTR_1g072530 | -0,03 | 0,08 | -0,02 | -0,04 |
| MTR_1g077640 | 0,35 | -0,50 | 0,06 | 0,09 |

**Table 4.**
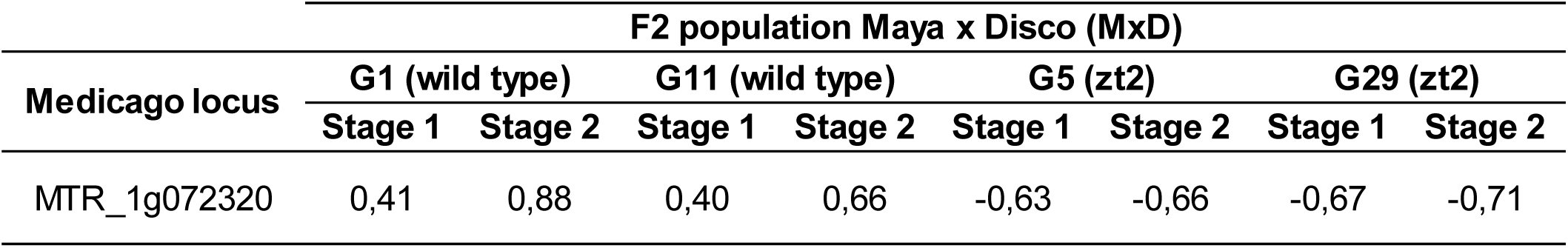
Validation of the candidate gene MTR_1g072320 in four F2 genotypes from the M x D population. Values indicate log average expression ratios of three biological replicates from each of the genotypes. Genotypes G1-G11 (wild type) and G5-G29 (mutant type). All the samples showed a statistically significant regulation (p < 0.001).

**Fig. 2.**
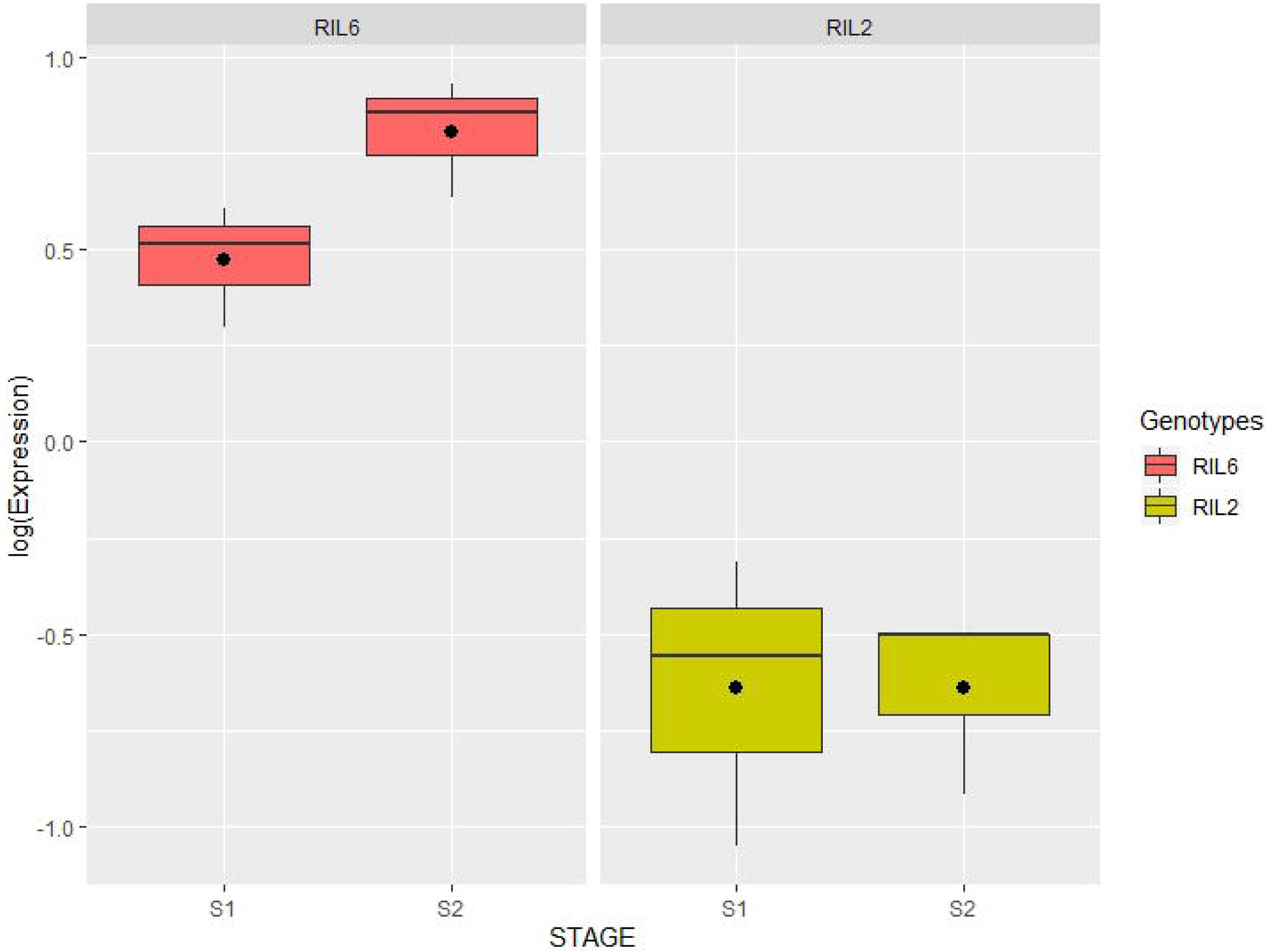
Boxplots showing the transcript levels of MTR_1g072320 in the RIL6 (pigmented flower) and the RIL2 (white flower) from cross V6 x zt2, in two developmental stages (S1: immature and S2: young flowers). The relative gene expression was log-transformed as detailed in Material and Methods. The expression values are shown as median (horizontal line), mean (asterisk), upper and lower quartiles (box) and range (whiskers).

**Fig. 3.**
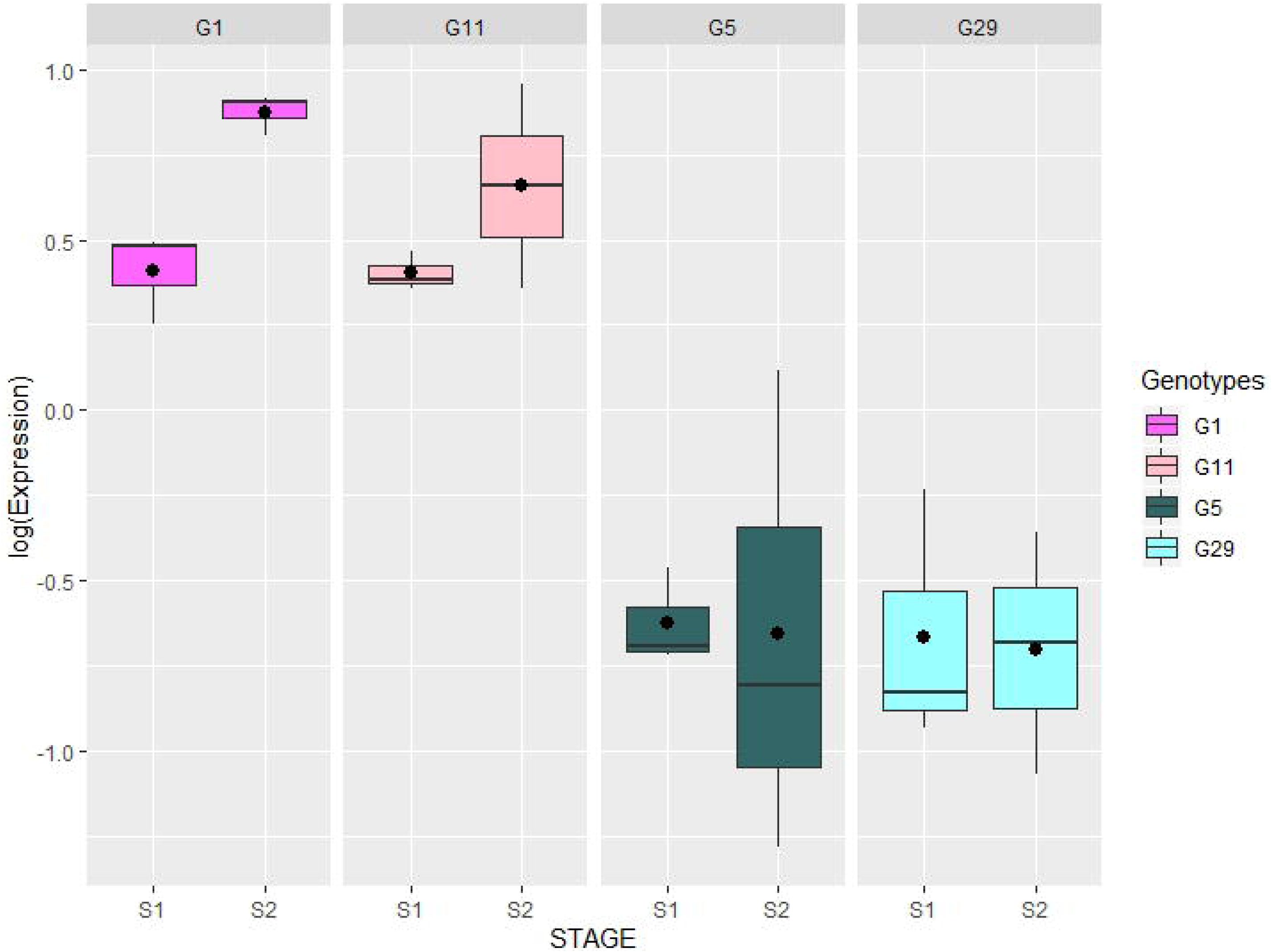
Validation of the MTR_1g072320 expression values in the F2 population M x D. Boxplots show the transcript levels in two individuals with pigmented flower (G1 and G11) and two white flowers (G5 and G29), in two developmental stages (S1: immature and S2: young flowers). The relative gene expression data was log-transformed as detailed in Material and Methods. The expression values are shown as median (horizontal line), mean (asterisk), upper and lower quartiles (box) and range (whiskers).

## Discussion

Faba bean is a valuable protein crop that could help the feed industry to shift towards a more sustainable raw material. Despite its prominent nutritional importance, faba bean also contain antinutrients, such as tannins, that reduce the bioavailability of proteins and minerals. Condensed tannins are found in coloured-flowering varieties of faba beans and are present in the seed coat. The zero-tannin (*zt*) trait is governed independently by two complementary recessive genes *zt1* and *zt2. TTG1* encodes the zt1 phenotypes^25–28^ but the gene underlying zt2 was still unknown. Therefore, identifying markers closely linked to *zt2* has been a key objective for increasing the accuracy and efficiency of selection and facilitate marker-assisted breeding of faba bean tannin-free cultivars.

Previous studies ascribed this locus to the distal part of chr. III^26,28,29^. The last authors also reported a KASP marker (SNP marker Vf_Mt7g100500_001), 10.5 cM far from flower colour, as a successful marker to discriminate low tannin plants although the closest flanking marker (Vf_Mt1g072740_001), 3 cM apart, failed to distinguish contrasted genotypes in the validation panel^29^. Contradicting the previous report, in this study Vf_Mt7g100500 revealed the highest number of misclassified individuals in any of the faba bean populations assayed preventing it use in marker assisted breeding.

To identify candidate transcription factors that might be responsible for the unpigmented *zt2* genotypes, marker enrichment of the target region was undertaken. Thus, a set of 23 MYB, bHLH and WD40 present in the collinear *M. truncatula* region where for the first time assayed in three different faba bean populations. Not all markers are applicable across populations due to lack of polymorphism. Multiple mapping populations of diverse genetic backgrounds are helpful for accurate fine-mapping and ultimately, for target gene identification. To maximize the genetic diversity, we have analyzed three faba bean segregating populations that increased the level of polymorphisms and narrowed down the search of potential candidates to a much shorter segment of the *M. truncatula* genome. The analysis revealed Vf_Mt1g072320, encoding a TT8 transcription factor as the candidate gene controlling zero tannin zt2 individuals.

In Arabidopsis, *TT8* (encoding a bHLH domain transcription factor), *TTG1* (encoding a WDR protein) and *TT2* (encoding a R2R3 MYB domain protein) have been identified as key determinants for the proanthocyanidin accumulation in developing seeds^42,43^. This MYB–bHLH–WD40 (MBW) complex TT2–TT8–TTG1 activates the transcription of structural genes in the anthocyanin pathway and regulates the accumulation of condensed tannins in the *Arabidopsis* seeds coat^19^. In Medicago, both WD40 repeats and bHLH transcription factors (MtTT8) are involved in the regulation of anthocyanin and proanthocyanidin biosynthesis^42,43^. In pea, the absence of pigmentation in the flower is the result of mutations in loci *A* and *A2* encoding a bHLH transcription factor and a WD40 protein, respectively^21^. A single recessive gene (*tan*) that encodes a bHLH transcription factor, homologous to the *A* gene in pea^22^ controls white flower in lentils, while in faba bean a WD40 transcription factor (VfTTG1) has been reported as the gene encoding the zt1 phenotypes^26,28,29^.

The conservation of the MBW complex controlling anthocyanin pigmentation in Eudicots^44^ suggests that other MYB or bHLH transcription factor should control *zt2* gene in faba bean. In this study the MtTT8 (Mt1g072320), ortholog to the genes *A* in pea and *tan* in lentil, was assayed as a promising candidate gene determining low tannin content in faba bean. Vf_Mt1g072320 (VfTT8), was polymorphic in the three faba bean populations and strongly cosegregates with the white flower lines, thus supporting the identity of the candidate. These findings were further confirmed by qPCR analyses in different populations and genetic background. In all cases, the VfTT8 expression level was very low in white flowered types, in contrast to the high expression in wild genotypes.

From the data presented here, we can conclude that VfTT8 is the gene responsible for the zt2 phenotypes in faba bean. Although further analysis is necessary to elucidate the mechanistic basis for the observed down-regulation of this gene, our results open the door to the development of an ultimate diagnostic marker based on the allelic variant causing the phenotypic effect. This study increases our understanding of the regulatory mechanisms underlying tannin biosynthesis in this crop and will likely favour the upcoming development of a fast and reliable tool for the generation of value-added faba bean cultivars with optimized flavonoid content.

## Acknowledgements

This research was supported by funding from the EU project EUCLEG (n°727312-2), as well as from the projects RTA2017-00041 and PP.AVA.AVA2019.030, all of them co-financed by ERDF and the latter by the regional government of Andalucía. The authors would like to thank Dr. A Di Pietro for his careful reading of the manuscript and helpful comments and suggestions, Dr. I Casimiro-Soriguer for her valuable assistance in the statistical analysis and Dr. C De Miguel Rojas for her valuable laboratory assistance.

## Authors contributions Statement

NG, CMA and AMT conceived and designed the study and lab experiments. NG performed the experiments, the analysis and the interpretation of the data. NG, CMA and AMT wrote the manuscript.

## Data Available

The datasets generated and analyzed during the current study are available from the corresponding author on request.

